# Murine SEC24D Can Substitute Functionally for SEC24C *in vivo*

**DOI:** 10.1101/284398

**Authors:** Elizabeth J. Adams, Rami Khoriaty, Anna Kiseleva, Audrey C. A. Cleuren, Kärt Tomberg, Martijn A. van der Ent, Peter Gergics, K. Sue O’Shea, Thomas L. Saunders, David Ginsburg

**Affiliations:** From the Life Sciences Institute, University of Michigan, Ann Arbor, MI 48109; Program in Cellular and Molecular Biology, University of Michigan, Ann Arbor, MI 48109; Department of Internal Medicine, University of Michigan, Ann Arbor, MI 48109; Departement of Human Genetics, University of Michigan, Ann Arbor, MI 48109; Department of Cell and Developmental Biology, University of Michigan, Ann Arbor, MI 48109; Department of Pediatrics, University of Michigan, Ann Arbor, MI 48109; Howard Hughes Medical Institute, University of Michigan, Ann Arbor, MI 48109

## Abstract

The COPII component SEC24 mediates the recruitment of transmembrane cargoes or cargo adaptors into newly forming COPII vesicles on the ER membrane. Mammalian genomes encode four *Sec24* paralogs (*Sec24a-d)*, with two subfamilies based on sequence homology (SEC24A/B and C/D), though little is known about their comparative functions and cargo-specificities. Complete deficiency for *Sec24d* results in very early embryonic lethality in mice (before the 8 cell stage), with later embryonic lethality (E 7.5) observed in *Sec24c* null mice. To test the potential overlap in function between SEC24C/D, we employed dual recombinase mediated cassette exchange to generate a *Sec24c*^*c-d*^ allele, in which the C-terminal 90% of SEC24C has been replaced by SEC24D coding sequence. In contrast to the embryonic lethality at E7.5 of SEC24C-deficiency, *Sec24c*^*c-d/c-d*^ pups survive to term, though dying shortly after birth. *Sec24c*^*c-d/c-d*^ pups are smaller in size, but exhibit no obvious developmental abnormality. These results suggest that tissue-specific and/or stage-specific expression of the *Sec24c/d* genes rather than differences in cargo function explain the early embryonic requirements for SEC24C and SEC24D.

## INTRODUCTION

In eukaryotic cells, most proteins destined for export from the cell, to the cell surface, or to various intracellular storage compartments must traverse the secretory pathway before reaching their final intracellular or extracellular destinations (1,2). The first step of this fundamental process is the concentration and packaging of newly synthesized proteins into vesicles on the surface of the ER at specific ER exit sites (3). At these sites, cytosolic components assemble to form the COPII complex (4,5), a protein coat that generates membrane curvature and promotes the recruitment of cargo proteins into a nascent COPII bud (6,7). Central to this process is SEC24, the COPII protein component primarily responsible for interaction between transmembrane cargoes (or cargo-bound receptors) and the coat (8). SEC24 forms a complex with SEC23 in the cytosol, and the SEC23/SEC24 heterodimer is drawn to ER exits sites upon activation of the GTPase SAR1(9) by its cognate ER membrane bound GEF, SEC12 (10). Once recruited to the ER membrane, SEC24 interacts with ER exit signals on the cytoplasmic tail of protein cargoes via cargo recognition sites and facilitate vesicle formation.

Mammalian genomes encode four SEC24 paralogs (*Sec24a-d*), with each containing several highly conserved C-terminal domains and a hypervariable N-terminal segment comprising approximately one-third of the protein sequence. Based on sequence identity, the four mammalian SEC24s can be further sub-divided into two subgroups, SEC24A/B and SEC24C/D (11). Murine SEC24A and B share 58% amino acid identity and murine SEC24C and D 60% identity, with only 25% amino acid sequence identity between the two subgroups (12), suggesting both ancient and more recent gene duplications. Mice with deficiency in the individual SEC24 paralogs exhibit a wide range of phenotypes. SEC24A-deficient mice present with markedly reduced plasma cholesterol as a result of impaired secretion of PCSK9, a regulatory protein that mediates LDL receptor degradation (13). SEC24B-deficient mice exhibit late embryonic lethality ∼E18.5 due to neural tube closure defects resulting from reduced trafficking of the planar-cell-polarity protein VANGL2 (14). Loss of SEC24C in the mouse results in embryonic lethality at ∼E7.5 (12), and SEC24D deficiency in mice results in embryonic death at or before the 8-cell stage (15), whereas mutations in human SEC24D have been reported in patients with certain skeletal disorders (16),

The expansion of the number of COPII paralogs over evolutionary time suggests a divergence in cargo recognition function, with the disparate deficient mouse phenotypes resulting from paralog specific protein interactions with a subset of cargo molecules, or other functions beyond cargo recognition. Consistent with this model, all four mammalian *Sec24s* are broadly expressed (15,17,18). Nonetheless, subtle differences in the developmental timing and/or tissue-specific patterns of expression as the explanation for the unique phenotypes associated with deficiency for each SEC24 paralog cannot be excluded.

To test the functional overlap between SEC24C/D *in vivo*, we employed dual recombinase mediated cassette exchange (dRMCE) (19) to knock-in the C-terminal 90% of the coding sequence for SEC24D in place of the corresponding SEC24C coding sequence, at the *Sec24c* locus. Surprisingly, these SEC24D sequences can largely substitute for SEC24C, rescuing the early embryonic lethality previously observed in SEC24C-deficient mice. In contrast, these SEC24D sequences expressed in the context of the *Sec24c* gene fail to rescue SEC24D-deficiency. These results demonstrate a high degree of functional overlap between the SEC24C/D proteins and suggest that the deficiency phenotypes for each paralog are determined largely by tissue or developmental-timing specific differences in their gene expression programs.

## RESULTS

### Identification of a dRMCE targeted ES cell clone

Direct microinjection of 102 zygotes generated from a *Sec24c*^*+/-*^ X *Sec24c*^*+/+*^ cross with pDIRE and the *Sec24c-d* replacement construct (Figure 1) failed to generate any progeny mice with the correctly targeted event; 65 were heterozygous for the *Sec24c*^-^ allele containing the FRT and loxP sites required for dRMCE (19) and 47 were wild type. While no targeted insertions were observed, 20/102 (20%) pups carried random insertions of the dRMCE replacement vector.

**Figure 1:**
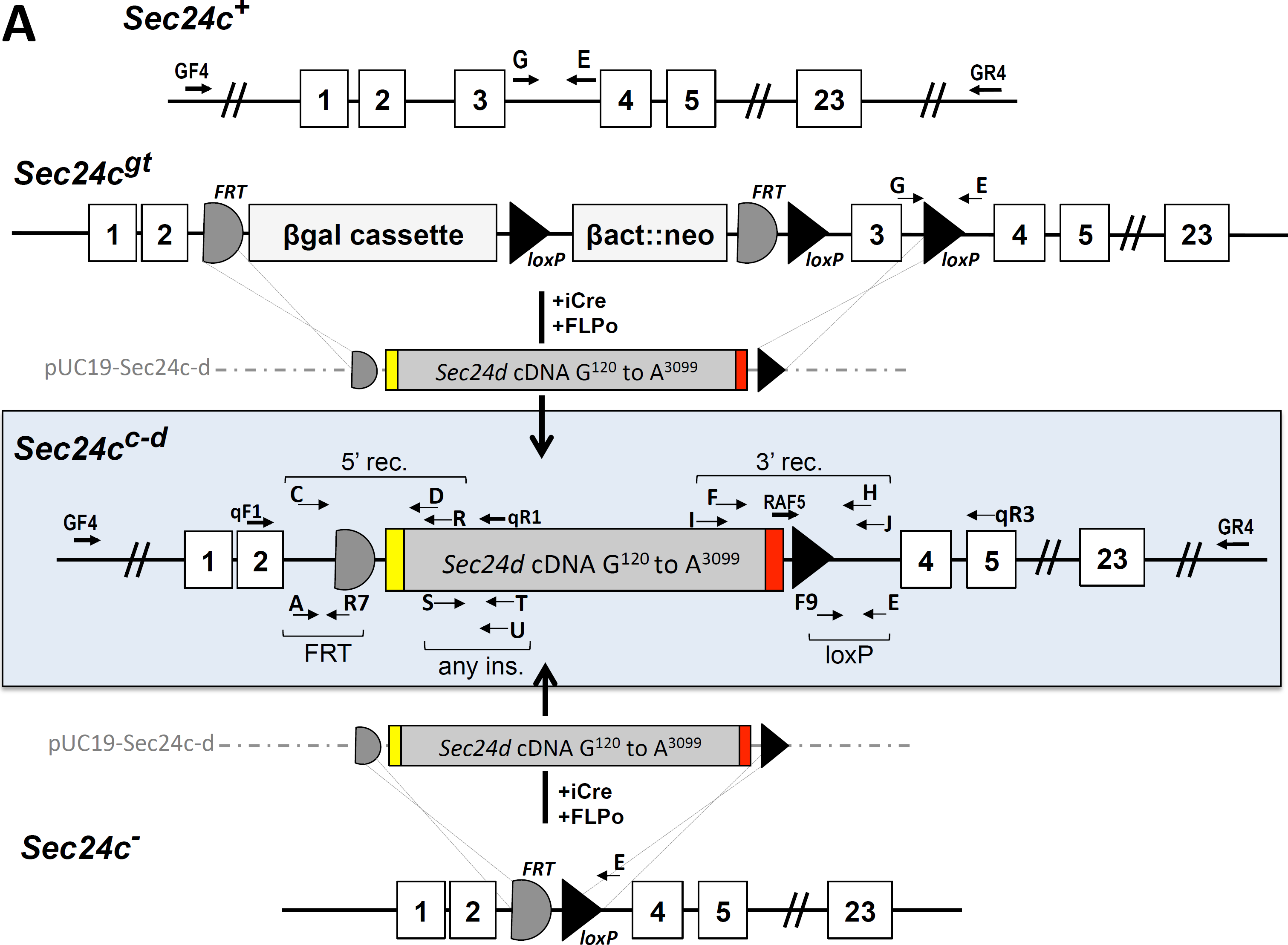
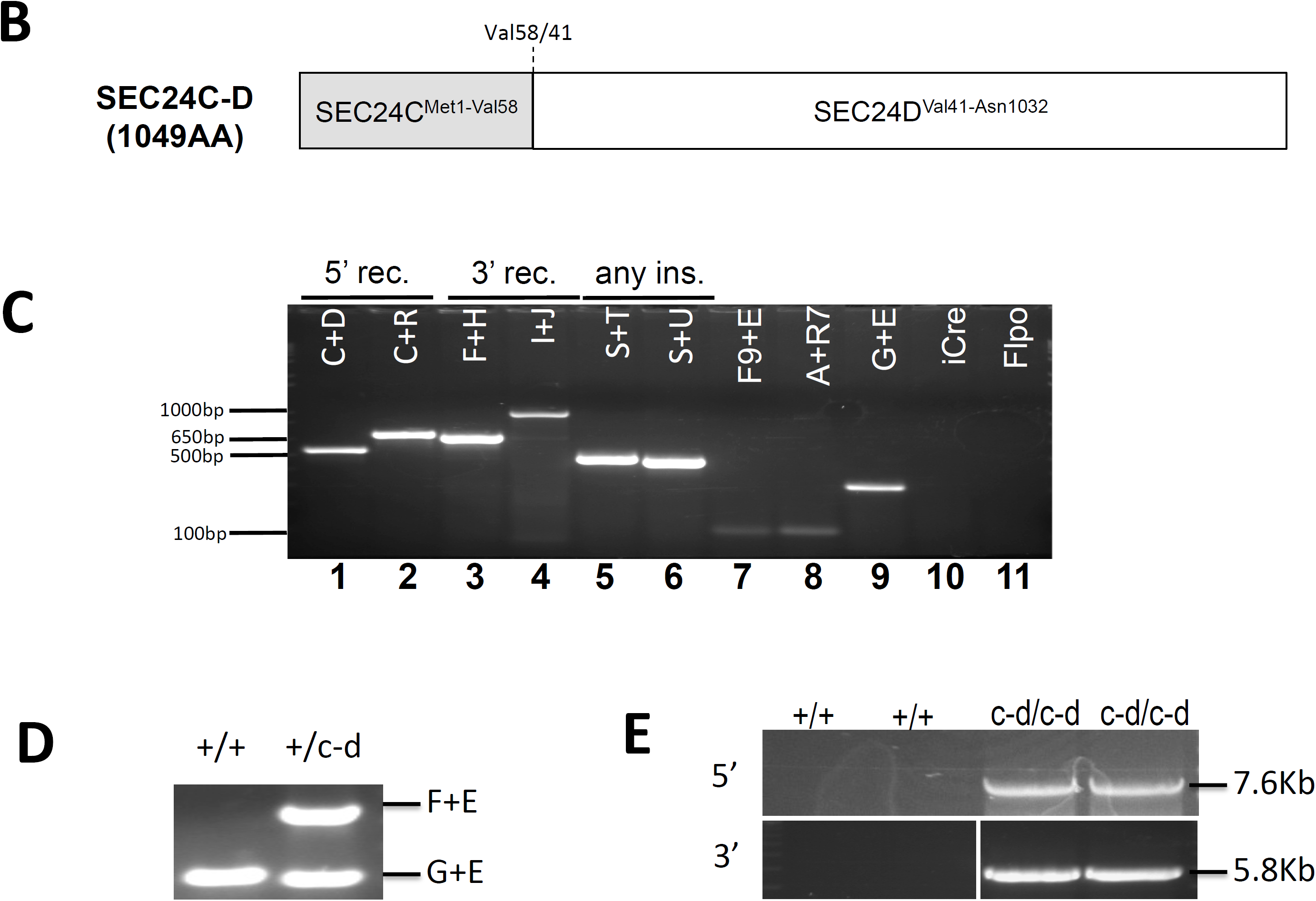
Design and generation of the chimeric *Sec24c^c-d^* allele. (A) Schematic representation of dRMCE to generate the *Sec24c*^*c-d*^ allele. The replacement vector pUC19-Sec24c-d contains the *Sec24c* intron 2 splice acceptor (yellow), the *Sec24d* coding sequence beginning with G^120^ (gray), and a stop codon followed by a poly A signal sequence (red). Arrows represent primers used for genotyping, long-range PCR, and RT-PCR (for sequences, see Table 4). (B) The SEC24C-D fusion protein encoded by the dRMCE generated *Sec24c*^*c-d*^ allele, which contains the first 57 amino acids of SEC24C followed by the SEC24D sequence corresponding to the remaining SEC24C sequence. Val58/41 (Val58 in SEC24C, Val41 in SEC24D) indicates the junction point for this chimeric protein (see also Supplemental Figure 1). (C) PCR results for dRMCE subclone 12275. Primer combinations are indicated at the top of each lane. Correct targeting was observed for the 5’ recombination (5’ rec) site (lanes 1 and 2) and the 3’ recombination (3’ rec) site (lanes 3 and 4). Additionally, the presence of the *Sec24d* cDNA (“any ins.”) (lanes 5,6), and the loxP and FRT sites (103bp and 106bp products in lanes 7,8, respectively) was confirmed. The signal in lane 9 is due to the *Sec24c*^*+*^allele, confirming that ESC clone 12275 is heterozygous for the *Sec24c*^*c-d*^ allele. Clone 12275 does not carry any random insertions of pCAGGS-iCre (lane 10) or pCAGGS-Flpo (lane 11). (D) A genotyping PCR assay to distinguish between the wild type and *Sec24c*^*c-d*^ allele using primers E, F, and G. (E) Long range PCR confirms correct targeting. Primers GF4+U were used to amplify the 5’ arm resulting in a 7.6kb product, and primers F and GR4 were used to amplify the 3’ arm to yield a 5.8kb product. Primers were located outside the homology arms (GF4 and GR4) and within the *Sec24d* cDNA (F and U). As expected, neither set of primers yields a band from the *Sec24c*^*wt*^ allele.

A screen of 288 ES cell clones co-electroporated with pDIRE and the *Sec24c-d* targeting construct also failed to yield any properly targeted colonies, though 18 random insertions of the *Sec24c-d* construct (6.25%, Table 1) were observed. However, analysis of a second set of 288 ES cell clones transfected with alternative Cre and FLP expression vectors identified a single clone carrying a potential targeted insertion of the dRMCE replacement construct into the *Sec24c* locus. A single round of subcloning generated pure clonal ES cell populations carrying the *Sec24c*^*c-d*^ allele.

**Table 1:**
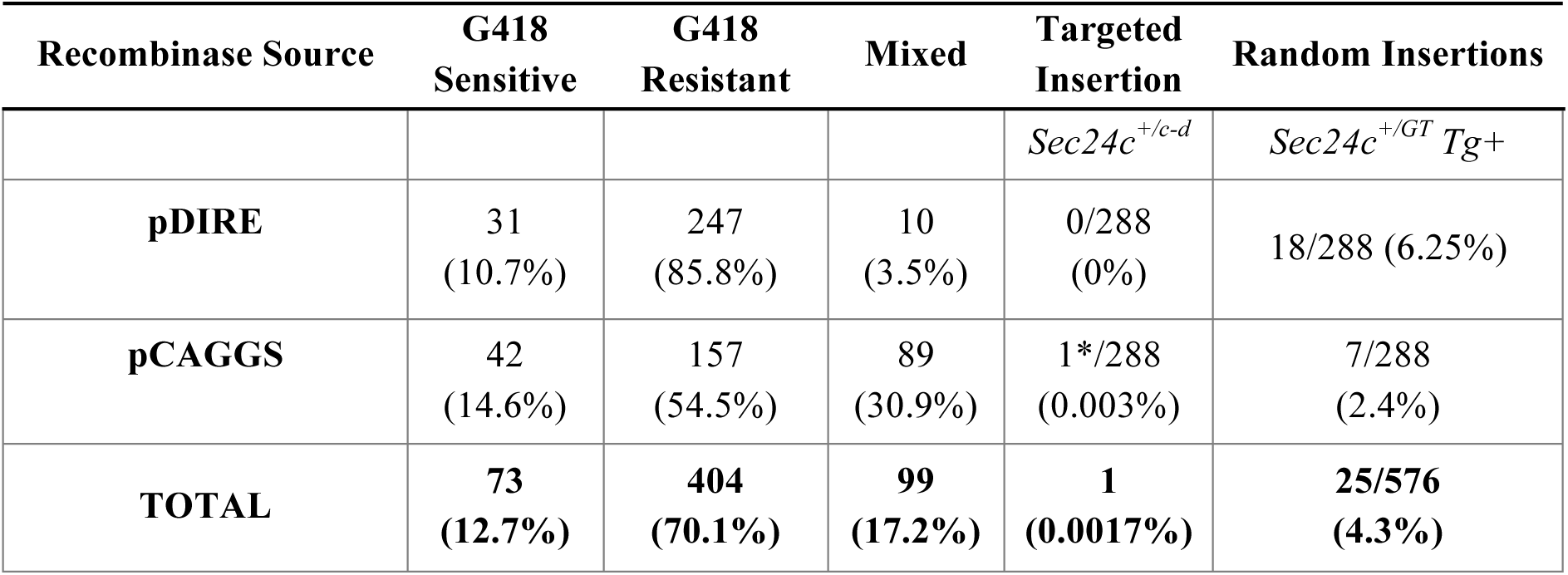
Summary of embryonic stem cell co-electroporation results. Results of co-electroporation of *Sec24c*^*+/GT*^ embryonic stem cells (ESCs) with pUC19-Sec24c-d and either pDIRE or pCAGGS-iCre and pCAGGS-Flpo. G418 resistance should indicate absence of recombination to remove the neomycin cassette present in the parental *Sec24c*^*GT*^ allele. The targeted clone identified (*=6-H9) contained a mixed population of G418 sensitive and resistant ESCs.

### No phenotypic abnormalities observed in *Sec24c^+/c-d^* mice

One out of the three ES cell clones microinjected achieved germline transmission, and mice carrying the *Sec24c*^*c-d*^ allele generated. The expected Mendelian ratio of *Sec24c*^*+/c-d*^ mice was observed in N2 progeny of backcrosses to C57BL/6J mice (*p* >0.38, Table 2A). *Sec24c*^*+/c-d*^ mice were indistinguishable from their wild type littermates, exhibiting normal fertility, and no gross abnormalities on standard autopsy examination. There were also no differences in body weight at 4 and 6 weeks of age (Figure 2). *Sec24c*^*+/c-d*^ *Sec24d*^*+/GT*^ mice are also viable and healthy and observed in the expected numbers (*p>*0.72, Table 2B-C). While only a small number of wild type offspring were maintained beyond 100 days of age, there was no significant difference in lifespan between *Sec24c*^*+/c-d*^ mice (n=93) or *Sec24c*^*+/c-d*^ *Sec24d*^*+/GT*^ mice (n= 24) compared to controls.

**Table 2:**
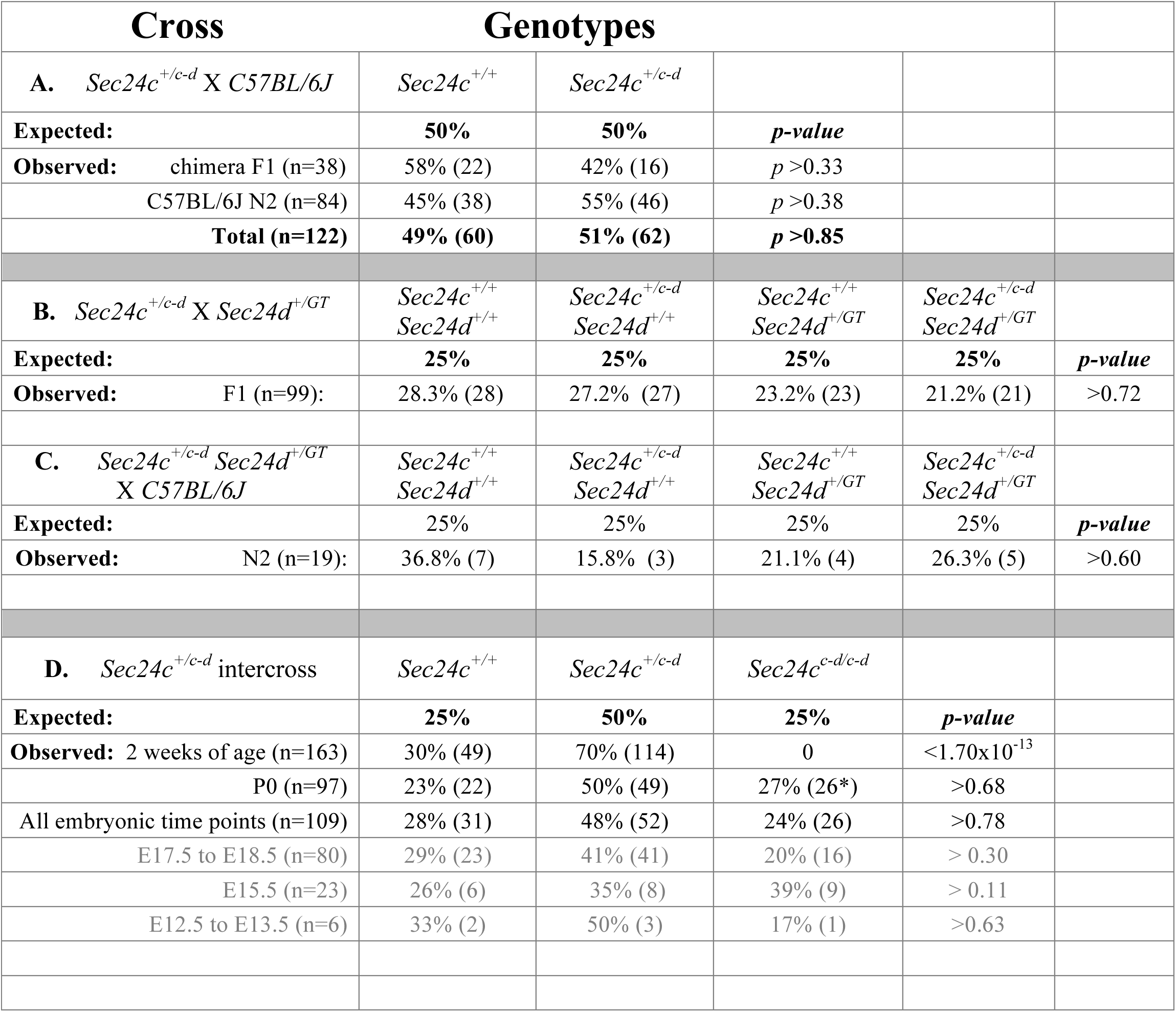
Results of *Sec24c^+/c-d^* backcrosses and intercrosses in this study. Distribution of progeny from (A) *Sec24c*^*+/c-d*^ backcrosses (B) *Sec24c*^*+/c-d*^ x *Sec24d*^*+/GT*^ intercrosses, (C) *Sec24c*^*+/c-d*^ *Sec24d*^*+/GT*^ backcrosses and (D) *Sec24c*^*+/c-d*^ intercrosses. Genotypes shown for chimera F1 only include those of chimera/ B6(Cg)-Tyr^c-2J^/J F1 progeny with black coat color. All *Sec24d*^*+/GT*^ mice used in this study were ≥N18 on C57BL/6J background. For intercross data, *p-*values are calculated based on “others vs. rescue” genotypes (0.75:0.25). *all Sec24c^c-d/c-d^ mice were dead at P0. The E15.5 time point in (D) contains genotypes for a single *Sec24c*^*+/c-d*^ resorbed embryo and two *Sec24c*^*c-d/c-d*^ absorbed embryos. All observed numbers are listed in parentheses.

**Figure 2:**
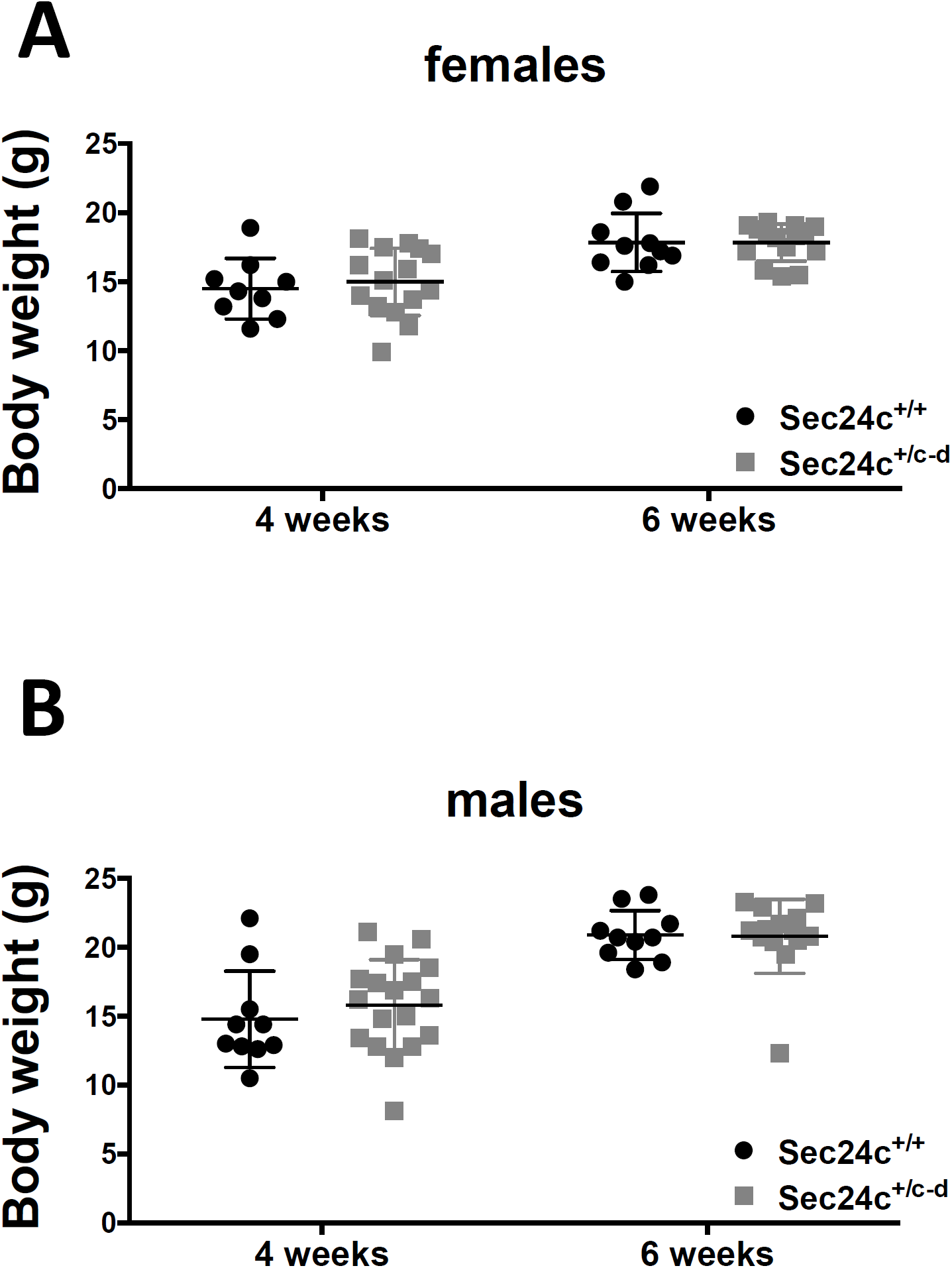
Body weights of *Sec24c^+/c-d^* mice. Mice weighed at 4 and 6 weeks of age for both *Sec24c*^*+/c-d*^ mice and wild type littermate controls show no significant difference between groups (p≥0.05). Error bars represent the standard deviation.

### The *Sec24c^c-d^* allele rescues *Sec24c^-/-^* mice from embryonic lethality

Table 2D shows the distribution of offspring from *Sec24c*^*+/c-d*^ intercrosses. No *Sec24c*^*c-d/c-d*^ mice were observed at weaning (0/163, *p*<1.7×10^−13^). Genotyping of 31 P0 progeny observed to die shortly after birth, all notably smaller and paler than their surviving littermates (Figure 3A), identified 26 as *Sec24c*^*c-d/c-d*^ (*p<*3.8×10^−14^). Taken together with the full set of progeny genotypes at P0 (n=97), the observed number of *Sec24c*^*c-d/c-d*^ offspring is consistent with the expected Mendelian ratios. *Sec24c*^*c-d/c-d*^ pups were 20-30% smaller by weight than their littermate controls at P0 (Figure 3B), were significantly shorter in crown-rump length (Figure 3C), and often exhibited a hunched appearance involving the shoulder girdle and trunk. Gross autopsy and histologic analyses (performed blinded to genotype) failed to identify any obvious abnormality to account for the neonatal lethality in *Sec24c*^*c-d/c-d*^ mice (Figure 4). Lungs of both heterozygous and wild type pups exhibited open alveoli lined by squamous epithelial cells (Figure 4B), while the alveoli of *Sec24c*^*c-d/c-d*^ neonates were open but had thickened walls (5/9) or were uninflated (4/9), and lined by columnar epithelia (Figure 4B).

**Figure 3:**
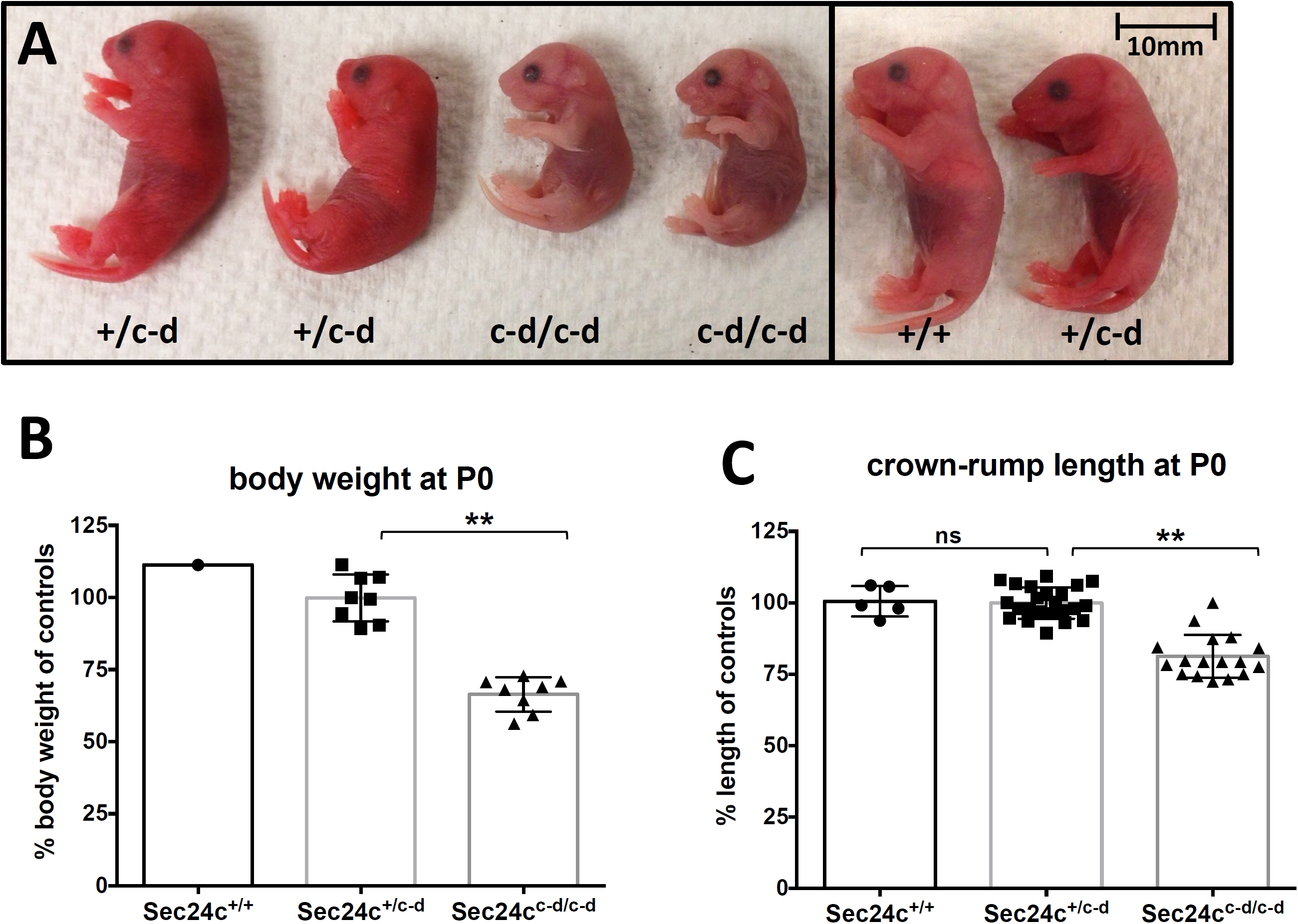
Phenotypic analysis of *Sec24c^+/c-d^* intercross progeny at P0. (A) Side views of P0 pups from *Sec24c*^*+/c-d*^ intercross taken shortly after birth. *Sec24c*^*c-d/c-d*^ pups are much paler and smaller than their littermate controls, exhibited little spontaneous movement, and died within minutes of birth. (B) Body weight measurements at P0, normalized to average weight of controls (*Sec24c*^*+/+*^ and *Sec24c*^*+/c-d*^) within the same litter. (C) Crown to rump length measurements at P0, normalized to average length of controls (*Sec24c*^*+/+*^ and *Sec24c*^*+/c-d*^) within the same litter. (**)= *p*<0.0001, (ns)= *p>*0.05. Error bars represent the standard deviation.

**Figure 4:**
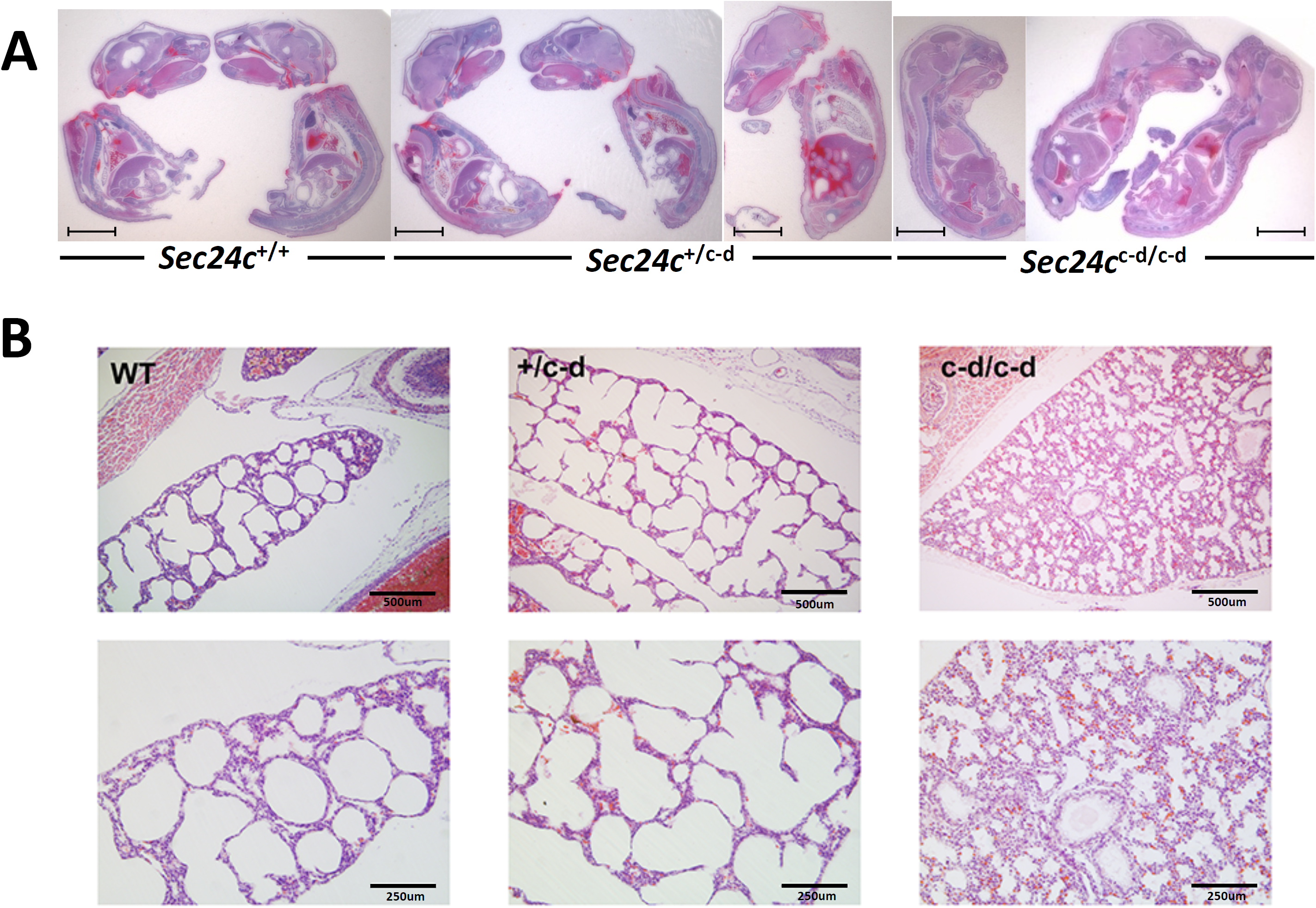
Histological assessment of *Sec24c^+/c-d^* intercross progeny at P0. (A) H&E stained longitudinal sections of P0 pups from *Sec24c*^*+/c-d*^ intercross taken shortly after birth. Scale bars = 5mm. (B) Low and higher magnification views of H&E stained sections through the lung of P0 mice collected shortly after birth. Analysis did not identify gross alterations in the morphology of the respiratory tree, although alveoli from *Sec24c*^*c-d/c-d*^ pups were often uninflated and lined by columnar epithelium compared to the squamous epithelium lining wild type and *Sec24c*^*+/c-d*^ alveoli. Lungs were fixed and sectioned in the context of the whole pup. **Scale bars = 500 microns.**

However, this lung pathology is unlikely to account for the neonatal death, as 7/9 neonates exhibit no breathing movements at delivery. At earlier embryonic time points (E12.5 to E17.5), viable *Sec24c*^*c-d/c-d*^ embryos were observed at the expected ratios (*p>*0.78), and though clearly smaller than their littermate controls, no other histopathological differences were identified at E15.5 or E17.5 upon observation by an independent reviewer blinded to sample genotypes (Figure 5). Thus, expression of SEC24C-D in place of SEC24C is sufficient to support development to term, but not survival past birth.

**Figure 5:**
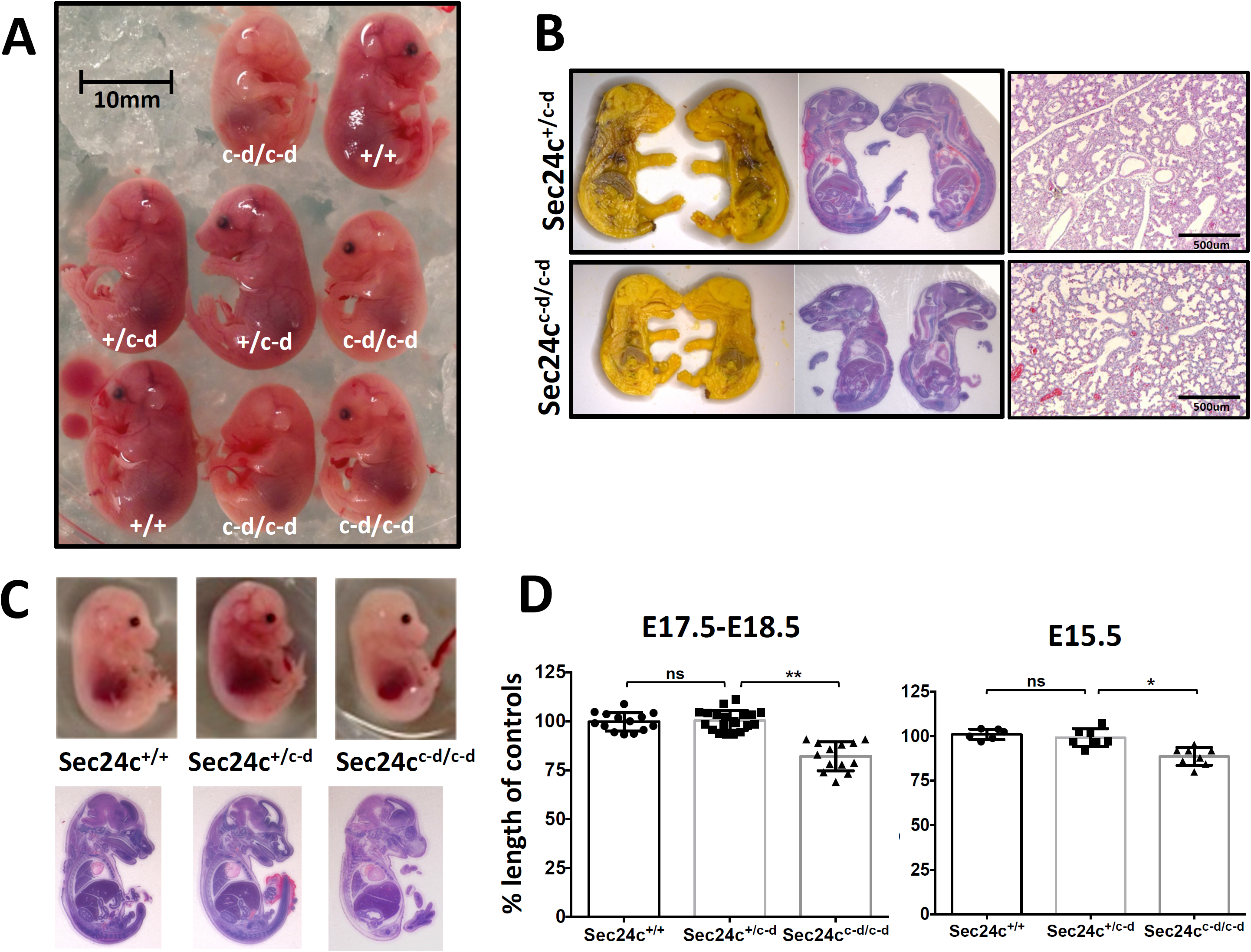
Phenotypic assessment of *Sec24c^+/c-d^* intercross embryos. (A) Side views of embryos from *Sec24c*^*+/c-d*^ intercross at E17.5. (B) Bouin’s solution fixed E18.5 embryos sectioned at the midline (left) and stained with H&E (center). H&E of 4% PFA fixed lungs from E18.5 embryos are shown at right. (C) Whole mount and H&E images of E15.5 embryos from *Sec24c*^*+/c-d*^ intercross. (D) Crown to rump length measurements at E17.5-18.5 and E15.5, normalized to average length of controls (*Sec24c*^*+/+*^ and *Sec24c*^*+/c-d*^) within the same litter. (**)= *p*<0.0001, (*)= *p<*0.002 (ns)= *p>*0.05. Error bars represent the standard deviation.

### The *Sec24c^c-d^* allele fails to complement disruption of Sec24d

Table 3 shows the results of intercrosses between *Sec24c*^*+/c-d*^ and *Sec24d*^*+/GT*^ (15) mice. No *Sec24c*^*+/c-d*^ *Sec24d*^*GT/GT*^ progeny were observed at embryonic time points from the blastocyst stage to birth (n=94, *p<*7.56×10^−5^) or at 2 weeks of age (n=113, *p<*1.43×10^−5^), although a single *Sec24c*^*+/c-d*^ *Sec24d*^*GT/GT*^ embryo was detected at the 8-cell pearly morula stage. These results are indistinguishable from the pattern previously reported for *Sec24d*^*GT/GT*^ mice (15).

**Table 3:** Results of *Sec24c^+/c-d^ Sec24d^+/GT^* x *Sec24d^+/GT^* intercrosses. For intercross data, p-values are calculated based on “others versus rescue” genotypes (6/7: 1/7). *Not considered a “rescue” because *Sec24d*^*GT/GT*^ morulas have been reported previously (15).

| Possible progeny of test cross: | *Sec24c*<sup>+/+</sup><br>*Sec24d*<sup>+/+</sup> | *Sec24c*<sup>+/+</sup><br>*Sec24d*<sup>+/*GT*</sup> | *Sec24c*<sup>+/+</sup><br>*Sec24d*<sup>*GT/GT*</sup> | *Sec24c*<sup>+/*c-d*</sup><br>*Sec24d*<sup>+/+</sup> | *Sec24c*<sup>+/*c-d*</sup><br>*Sec24d*<sup>+/*GT*</sup> | *Sec24c*<sup>+/*c-d*</sup><br>*Sec24d*<sup>*GT/GT*</sup> |  |
| --- | --- | --- | --- | --- | --- | --- | --- |
| Expected ratio with rescue: | 14.3% (1/7) | 28.6% (2/7) | 0 (early lethal *) | 14.3% (1/7) | 28.6% (2/7) | 14.3% (1/7) |  |
| Observed: |  |  |  |  |  |  | *p-value* |
| 2-weeks of age (n=113) | 14% (16) | 38% (43) | 0% | 12% (13) | 36% (41) | 0% | 1.43 x10<sup>-5</sup> |
| blastocyst to E18.5 (n=94) | 11% (10) | 37% (35) | 1% (1) | 11% (10) | 40% (38) | 0% | 7.55 x10<sup>-5</sup> |
| E15.5 to E18.5 (n=25) | 12% (3) | 40% (10) | 0% | 8% (2) | 40% (10) | 0% | 0.041 |
| E12.5 (n=29) | 14% (4) | 24% (7) | 0% | 21% (6) | 41% (12) | 0% | 0.0279 |
| blastocyst (n=40) | 10% (4) | 43% (17) | 3% (1) | 5% (2) | 40% (16) | 0% | 0.009 |
| 8 cell morula (n=9) | 0% | 33% (3) | 0% | 22% (2) | 33% (3) | 11% (1*) | 0.875 |
| **Total (n=216)** | **12.0% (26)** | **37.5% (81)** | **0.5% (1)** | **11.6% (25)** | **38.0% (82)** | **0.5% (1\*)** | **7.89 x10<sup>-7</sup>** |

### Low level of splicing around the *Sec24c^c-d^* allele

RT-PCR analysis of RNA from both *Sec24c*^*+/c-d*^ and *Sec24c*^*c-d/c-d*^ embryos detected the expected mRNA resulting from splicing of *Sec24c* exon 2 to the *Sec24c*^*c-d*^ fusion (Figure 6, lanes 3-6). However, a low level of splicing around the dRMCE insertion (directly from *Sec24c* exon 2 to exon 4) was also observed in both *Sec24c*^*+/c-d*^ and *Sec24c*^*c-d/c-d*^ embryos. DNA sequencing analysis confirmed that this product contained the sequence of exons 2-4-5, consistent with a splicing event around the dRMCE insertion. The absence of exon 3 in this transcript (also seen in the *Sec24c*^*+/-*^ samples) results in a frame-shift and early termination codon, as previously described (12).

**Figure 6:**
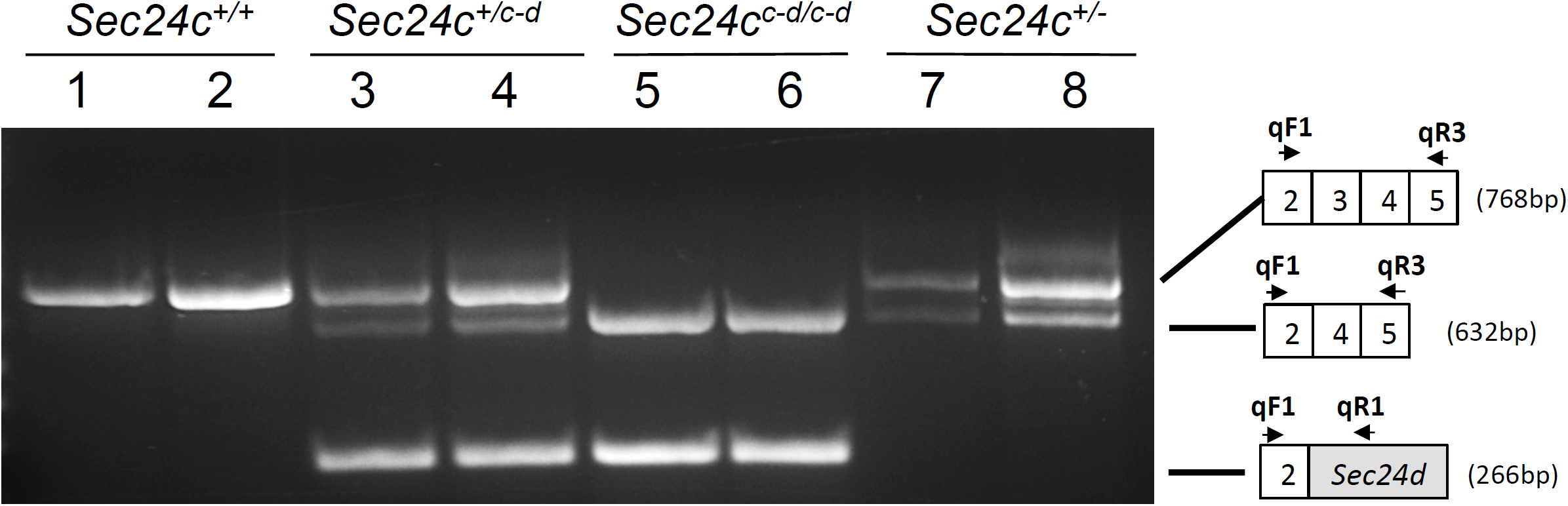
Analysis of *Sec24c^c-d^* allele splicing. A three-primer RT-PCR reaction using qF1, qR1 and qR3 *Sec24c*^*c-d*^ demonstrates the presence of *Sec24c*^*c-d*^ allele splice variants. Primers qF1 and qR3 (see schematic at the right) detected mRNA transcripts from both the wild type allele (exons 2-3-4-5) and the transcript skipping exon 3 (2-4-5) present in mice carrying the *Sec24c*^*c-d*^ allele, resulting from splicing around the dRMCE insertion (lanes 3-6), as well as mice carrying the *Sec24c*^-^ allele, in which exon 3 was also excised (lanes 7-8). Primers qF1 and qR1 detect the mRNA transcript of the *Sec24d* cDNA insert in *Sec24c*^*+/c-d*^ and *Sec24c*^*c-d/c-d*^ mice (lanes 3-6), and this transcript is absent in wild type littermates (lanes 1-2), or *Sec24c*^*+/-*^ mice (lanes 7-8).

## DISCUSSION

We show that SEC24D coding sequences inserted into the *Sec24c* locus (*Sec24c*^*c-d*^) largely rescue the early embryonic lethality observed in *Sec24c*^*-/-*^ mice (12). However, *Sec24c*^*c-d*^ is unable to substitute for SEC24D expression in *Sec24c*^*+/c-d*^ *Sec24d*^*GT/GT*^ mice. Our findings suggest that SEC24D can largely or completely substitute functionally for SEC24C. The incomplete rescue of SEC24C deficiency by the substituted SEC24D sequences could be due to imperfect interaction between the residual 57 amino acids of SEC24C retained at the N-terminus with the remaining 992 C-terminal amino acids of SEC24D. Alternatively, the targeting of *Sec24d* cDNA sequences into the *Sec24c* locus could have potentially disrupted regulatory sequences important for the control of *Sec24c* gene expression. Finally, it should be noted that there are two alternative splice forms of *Sec24c* with one form containing an additional 23 amino acid insertion, with the paralogous sequences absent from SEC24D (12). Thus, loss of a unique function conferred by this alternatively spliced form of SEC24C could also explain the perinatal lethality observed in *Sec24c*^*c-d/c-d*^ mice. Although our data are consistent with complete functional equivalence between the SEC24C and SEC24D proteins, we cannot exclude the possibility that subtle paralog-specific differences between the SEC24C and SEC24D proteins could account for the *Sec24c*^*c-d/c-d*^ phenotype.

Other examples of complementation by gene replacement include *Axin* and *Axin2/Conductin* (20), transcription factors Pax2 and Pax5 (21), cyclin D1 and cyclin E (22), En-1 and En-2 (23), N-*myc* and c-*myc* (24), Oxt2 and Oxt1 (25), and members of the Hox gene family, including *Hoxa3* and *Hoxd3* (26), all of which were carried out by traditional knock-in approaches with cDNA targeting constructs and homologus recombination. The proteins encoded by these genes are involved in key tissue-specific transcriptional and regulatory pathways, where such complementarity may be expected. In contrast, the COPII secretory pathway is an essential component of all eukaryotic cells, yet this study demonstrates that the SEC24 paralogs also have the capacity to largely substitute functionally for one another if placed in the proper expressional context. These paralogs strike a balance between functional redundancy and having completely unique functions, likely to allow higher organisms to efficiently meet the various demands secretory pathway in so many different cellular contexts. Nonetheless, the remarkable extension of survival from E7.5 to E18.5 and generally normal pattern of embryonic development in *Sec24c*^*c-d/c-d*^ mice demonstrate a high degree of functional overlap between SEC24C and D as well as the critical importance of spatial, temporal, and quantitative gene expression programs in determining the phenotypes of SEC24C and SEC24D deficiency.

Several secretory protein cargos have been shown to exhibit specificity for an individual SEC24 paralog, including the dependence of VANGL2 on SEC24B (14), the serotonin transporter (SERT) on SEC24C (27), and the GABA1 transporter on SEC24D (28). However, there is also evidence for significant overlap among the cargo repertoires of the mammalian *Sec24* paralogs, particularly within the subfamilies. Several cargo exit motifs are recognized by multiple SEC24 paralogs, including the DxE signal on VSV-G, and the IxM motif on syntaxin 5, both of which confer specificity for human SEC24A/B (29). The human transmembrane protein p24-p23 exhibits a preference for SEC24C or SEC24D and is thought to be a cargo receptor for GPI-anchored CD59, explaining the specificity of the latter for SEC24C/D (30). Similarly, PCSK9, which is dependent on SEC24A for ER export shows some overlap with SEC24B, both *in vivo* and *in vitro*, but none with SEC24C or D (13).

Taken together with our results, these findings suggest significant functional overlap within but not between the SEC24A/B and SEC24C/D subfamilies. This model is also consistent with the ∼55-60% sequence identity between SEC24A and B and between SEC24C and D, but only ∼25% between the A/B and C/D subfamilies. This high degree of functional overlap within SEC24 subfamilies also suggests that the deficiency phenotypes observed for loss of function for any of the SEC24 paralogs is due in large part to subtle differences in their finely tuned expression patterns, despite reports of generally ubiquitous expression for all 4 paralogs (12).

Humans with compound heterozygous mutations in SEC24D present with skeletal disorders such as Cole-Carpenter syndrome and severe osteogenesis imperfecta (16), with the medaka *vbi* (31) and the zebrafish *bulldog* (32) mutants exhibiting similar skeletal defects. These results have been interpreted as indicating a specific critical role for SEC24D in the secretion of extracellular matrix proteins (16,31,32). However, SEC24D-deficient mice exhibit very early embryonic lethality (15), at a time in development well before the establishment of the skeletal system. A similar discrepancy in phenotypes between mice and humans has been observed for SEC23B deficiency, which manifests as congenital dyserythropoietic anemia type II in humans (33), in contrast to perinatal lethality due to pancreatic disruption in mice (34,35). Our data suggest that evolutionary shifts in the expression programs for the *Sec24c* and *Sec24d* genes may explain these disparate phenotypes across vertebrate species, despite considerable overlap at the level of SEC24C/D protein function. Such changes in relative levels of gene expression could result in major differences in dependence on one or the other paralog across tissue types, even among closely related species.

## MATERIALS AND METHODS

### Cloning of *Sec24c-d* dRMCE construct pUC19-Sec24c-d

pUC19-Sec24c-d (Figure 1) was generated by assembling the Sec24c-d cassette (GenBank accession KP896524) which contains a FRT sequence, the endogenous *Sec24c* intron 2 splice acceptor sequence, a partial *Sec24d* coding sequence (from G^120^ to A^3099^ in the cDNA sequence, encoding the SEC24D sequence starting at Val41 (Supplemental Figure 1)) and the SV40 polyA sequence present in the *Sec24c*^*GT*^ allele (12). The entire cassette was inserted into pUC19 at the HindIII and EcoRI digests, and the integrity of the sequence was confirmed by DNA sequencing.

### Plasmid purification and microinjections

pDIRE, the plasmid directing dual expression of both iCre and FLPo (19) was obtained from Addgene (Plasmid 26745). Plasmid pCAGGS-FLPo was prepared by subcloning the FLPo-bovine growth hormone polyadeylation signal sequences from the pFLPo plasmid (36) into a pCAAGS promoter plasmid (37). Plasmid pCAAGS-iCre was prepared by adding the bovine growth hormone polyadenylation signal to iCre (kind gift of Rolf Sprengel) (38) and subcloning into a pCAGGS promoter plasmid. Plasmids pCAGGS-iCre and pCAGGS-FLPo contain the CAG promoter/enhancer, which drives recombinase expression of iCre or FLPo in fertilized mouse eggs (39), and were used as an alternative source of iCre and FLPo for some experiments, as noted. All plasmids, including the *Sec24c-d* replacement construct described below, were purified using the Machery-Nagel NucleoBond^®^ Xtra Maxi EF kit, per manufacture’s instructions. All microinjections were carried out at the University of Michigan Transgenic Animal Model Core. Co-injections of pUC19-Sec24c-d with pDIRE were performed on zygotes generated from the *in vitro* fertilization of C57BL/6J oocytes with sperm from *Sec24*^*+/-*^ male mice (12). For each microinjection, 5ng/μl of circular recombinase plasmid mixed with 5ng/μl of circular donor plasmid was administered, as described previously (40,41). Microinjected zygotes were then transferred to pseudopregnant foster mothers. Tail clips for genomic DNA isolation were obtained from pups at 2 weeks of age.

### Genotyping Assays

Genotypes of potential transgenic mice and ES cell clones were determined using a series of PCR reactions at the *Sec24c* locus. All genotyping primers used in this study are listed in Table 4 and those for the *Sec24c* locus are depicted in Figure 1, and expected band sizes are given in Table 5. Primers S, T, and U were used to amplify fragments of DNA unique to pUC19-Sec24c-d, which should detect targeted insertion at the *Sec24c* locus or random insertion elsewhere in the genome. Targeted insertion was detected using primer pairs flanking the FRT 5’ recombination site (primers C + primer D or R) and the loxP 3’ recombination site (primers I or F + primers H or J). Integration of the pCAGGS-iCre, pCAGGS-FLPo, or pDIRE was detected with primer sets iCre10F + iCre10R or FLPo8F + FLPo8R. Mice carrying the *Sec24d*^*gt*^ allele were genotyped as describe previously (15).

**Table 4:**
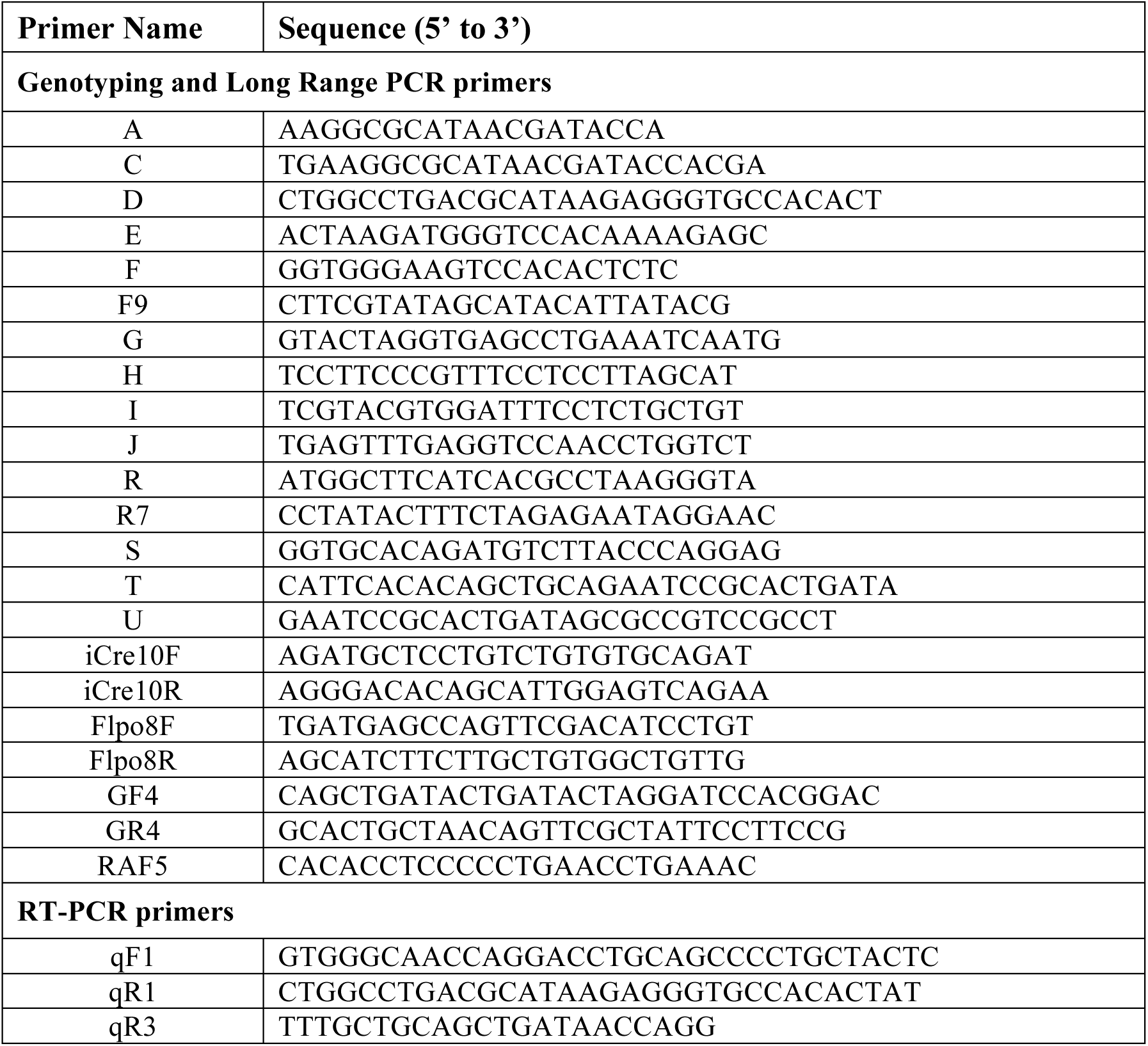
List of primers used in this study. All primer sequences listed 5’ to 3’.

**Table 5:** Expected PCR products from various Sec24c alleles. Primer pairs listed correspond to assays carried out either screening PCRs of ESC clones, genotyping of mice potentially carrying the *Sec24c*^*c-d*^ allele, and long range PCRs confirming the genomic location of the modified *Sec24c* locus. *632bp product resulting from splicing around the Sec24c-d insertion to exon 5.

| Primer Pairs | *Sec24c*<sup>+</sup> | *Sec24c*<sup>c-d</sup> | *Sec24c*<sup>gt</sup> | Type |
| --- | --- | --- | --- | --- |
| C+D | -- | 563 | -- | Screening PCR |
| C+R | -- | 727 | -- | Screening PCR |
| F+H | -- | 689 | -- | Screening PCR |
| I+J | -- | 993 | -- | Screening PCR |
| S+T | -- | 484 | -- | Screening PCR |
| S+U | -- | 468 | -- | Screening PCR |
| F9+E | -- | 103 | 103 | Screening PCR |
| A+R7 | -- | 106 | 106 | Screening PCR |
| G+E | 308 | -- | 308 | Genotyping |
| F+E | -- | 568 | -- | Genotyping |
| qF1+qR1 | -- | 266 | -- | RT-PCR |
| qF1+qR3 | 768 | 632* | -- | RT-PCR |
| GF4+U | -- | 7593 | -- | Long range PCR |
| RAF5+ GR4 | -- | 5798 | 6647 | Long range PCR |

### Transient electroporation of ES cells

ES cell clone EPD0241-2-A11 for *Sec24c*^*tm1a(EUCOMM)Wtsi*^ (*Sec24c*^*gt*^ allele,(12)) was expanded and co-electroporated with pUC19-Sec24c-d and either pDIRE or pCAGGS-iCre and pCAGGS-FLPo. ES culture conditions for JM8.N4 ES cells were as recommended at www.KOMP.org. Electroporation was carried out as previously described (19) with the exception that pUC19-Sec24c-d lacks a drug selection cassette that can be used to enrich for correct homologous recombination in mouse ES cells. After 1 week, individual ES cell colonies were plated in 96 well plates, 284 from cells transfected with pUC19-Sec24c-d and pDIRE, and 284 from cells transfected with pUC19-Sec24c-d, pCAGGS-iCre, and pCAGGS-FLPo. Cells were expanded and plated in triplicate for frozen stocks, DNA analysis, and G418 screening. To test for G418 sensitivity, cells were fed with G418 containing media for 1 week, and then fixed, stained, and evaluated for growth. Genomic DNA was prepared from each ES cell clone as previously described (42) and resuspended in TE.

### Subcloning of ES cells

PCR analysis demonstrated that clone 6-H9 contained a mixed population of cells, some of which were properly targeted with recombinations occurring at the outermost FRT and loxP sites, and others that still contained the parental *Sec24c*^*gt*^ allele, consistent with the observed mixed resistance to G418 (Table 1). One round of subcloning produced six different subclones consisting of a pure population of properly targeted ES cells, each with one wild type *Sec24c* allele and one correctly targeted *Sec24c*^*c-d*^ allele (Figure 1C) and none carrying random insertions of either pCAGGS-iCre or pCAGGS-FLPo.

### Generation of *Sec24c^+/c-d^* mice

Three correctly targeted subclones of 6-H9 were used to generate *Sec24c*^*+/c-d*^ mice (Figure 1D). ES cell clones were cultured as described previously (43) and expanded for microinjection. ES cell mouse chimeras were generated by microinjecting C57BL/6N ES cells into albino C57BL6 blastocysts as described (44) and then bred to B6(Cg)-Tyr^c-2J^/J (JAX stock #000058) to achieve germ-line transmission. ES-cell-derived F1 black progeny were genotyped using primers G, E, and F (Figure 1D). The *Sec24c*^*c-d*^ allele was maintained on the C57BL/6J background by continuous backcrosses to C57BL/6J mice. Initial generations were also genotyped to remove any potential iCre and FLPo insertions.

### Long-Range PCR

The integrity of the *Sec24c* locus with the newly inserted SEC24D sequence was confirmed by long-range PCR (Figure 1E). Genomic DNA from *Sec24c*^*+/+*^ and *Sec24c*^*+/c-d*^ mice were used as templates for a long-range PCR spanning the original arms of homology used for construction of the *Sec24c*^*gt*^ allele (12). Primers used for long range-PCR are depicted in Figure 1 and listed in Table 4. PCR was carried out using Phusion Hot Start II DNA Polymerase (Thermo Scientific), and products were fractionated on a 0.8% agarose gel.

### RT-PCR

Total RNA was isolated from a tail clip of *Sec24c*^*+/+*^, *Sec24c*^*+/c-d*^, and *Sec24c*^*c-d/c-d*^ embryos and liver biopsies from *Sec24c*^*+/-*^ mice using the RNAeasy kit (Qiagen) per manufacturer’s instructions, with the optional DNaseI digest step included. cDNA synthesis and PCR were carried out in one reaction using SuperScript® III One-Step RT-PCR System with Platinum®*Taq* (Invitrogen) following manufacturer’s instructions. Primers used for RT-PCR are depicted in Figure 6 and listed in Table 4.

### Timed Matings

Timed matings were carried out by intercrossing *Sec24c*^*+/c-d*^ mice. Embryos were harvested at designated time point for genotyping and histological analysis. Genotyping was performed on genomic DNA isolated from tail clip from mice >E12.5 or from yolk sacs and embryonic tissue from embryos < E12.5 days of age.

### Animal Care

All animal care and use complied with the Principles of Laboratory and Animal Care established by the National Society for Medical Research. The University of Michigan University Committee on Use and Care of Animals (UCUCA) approved all animal protocols in this study.

### Histology

Tissues, embryos and pups were fixed in Bouin’s solution (Sigma-Aldrich) at room temperature overnight, then transferred to 70% EtOH. Prior to embedding, fixed P0 pups and E17.5-E18.5 embryos were sectioned at the midline. Processing, embedding, sectioning and H&E staining were performed at the University of Michigan Microscopy and Image Analysis Laboratory.

### Embryonic Phenotyping

Crown to rump length measurements were obtained using a caliper on fresh specimens or by measuring the length of the embryo in a longitudinal section. Body weights were obtained immediately after birth. To account for normal variations in embryonic length and weight between litters, all measurements were normalized to the average length of *Sec24c*^*+/+*^ and *Sec24c*^*+/c-d*^ animals within a given litter (mean of controls = 100%). Values for each individual were then calculated based on that average for controls within the same litter.

### Statistical Analysis

To determine if there is a statistical deviation from the expected Mendelian ratios of genotypes from a given cross, the *p*-value reported is the χ^2^ value calculated using the observed ratio of genotypes compared to the expected ratio. All other *p-*values were calculated using Student’s unpaired t-test.

## Acknowledgements

We acknowledge Elizabeth Hughes, Corey Ziebell, Galina Gavrilina, Wanda Filipiak, Keith Childs, and Debora VanHeyningen for microinjections and preparation of ES cell-mouse chimeras from dRMCE ES clone 12275 and the Transgenic Animal Model Core of the University of Michigan Biomedical Research Core Facilities. Core support was provided by the University of Michigan Multipurpose Arthritis Center, NIH grant number AR20557 and the University of Michigan Cancer Center, NIH grant number CA46592. All histology work was performed in the Microscopy and Image-analysis Laboratory (MIL) at the University of Michigan, Biomedical Research Core Facilities (BRCF) with the assistance of Judy Poore. The MIL is a multi-user imaging facility supported by NIH-NCI, O’Brien Renal Center, UM Medical School, Endowment for the Basic Sciences (EBS), Department of Cell & Developmental Biology (CDB), and the University of Michigan. E.J.A was partially supported by the Cellular and Molecular Biology Training Grant (T32-GM007315). This work was supported by grants from the National Institutes of Health (PO1HL057346, RO1HL039693, and K08HL128794). R.K. is recipient of American Society of Hematology Scholar Award. D.G. is an investigator of the Howard Hughes Medical Institute.

